# Deconvoluting drug interactions based on *M. tuberculosis* physiologic processes: Transcriptional disaggregation of the BPaL regimen *in vivo*

**DOI:** 10.1101/2025.02.24.639926

**Authors:** Elizabeth A Wynn, Christian Dide-Agossou, Reem Al Mubarak, Karen Rossmassler, Jo Hendrix, Martin I Voskuil, Andrés Obregón-Henao, Michael A Lyons, Gregory T Robertson, Camille M Moore, Nicholas D Walter

**Affiliations:** Rocky Mountain Regional VA Medical Center, Aurora, CO, USA; Center for Genes, Environment and Health, National Jewish Health, Denver, CO, USA; Consortium for Applied Microbial Metrics, Aurora, CO, USA; Division of Pulmonary Sciences and Critical Care Medicine, University of Colorado Anschutz Medical Campus, Aurora, CO, USA; Linda Crnic Institute for Down Syndrome, University of Colorado Anschutz Medical Campus, Aurora, CO, USA; Department of Immunology and Microbiology, University of Colorado Anschutz Medical Campus, Aurora, Colorado, USA; Mycobacteria Research Laboratories, Department of Microbiology, Immunology, and Pathology, Colorado State University, Fort Collins, CO, USA; Department of Biostatistics and Informatics, University of Colorado Anschutz Medical Campus, Aurora, CO, USA

## Abstract

Identification of optimal antibiotic combination treatments for tuberculosis (TB) in preclinical studies is impeded by the limited information conventional pharmacodynamic (PD) markers provide about drug interactions. Measurement of individual drug activity based on colony forming units (CFU) does not reliably predict the activity of drug combinations, potentially because one drug may affect the physiology of *Mycobacterium tuberculosis* (*Mtb*) in a way that either favors or disfavors the activity of a second drug. SEARCH-TB is a novel candidate PD approach which uses targeted *in vivo* transcriptional profiling to evaluate the effects of drugs on *Mtb* physiology. To test the capacity of SEARCH-TB to elucidate drug interactions, we deconstructed the BPaL (bedaquiline, pretomanid, linezolid) regimen in the BALB/c high-dose aerosol mouse infection model, measuring the effect of 2, 7, and 14-day treatment with drugs in monotherapy, pairwise combinations, and as a 3-drug combination. Monotherapy rapidly induced drug-specific *Mtb* transcriptional responses by day 2 with continued evolution over 14 days. Bedaquiline dominated pairwise combinations with both pretomanid and linezolid. The pretomanid-linezolid combination gave a blended response, inducing transcriptional profiles “intermediate” between either drug. In the 3-drug BPaL regimen, the addition of both pretomanid and linezolid to bedaquiline yielded a greater transcriptional response than expected based on pairwise results. This work demonstrates that physiologic perturbations induced by a single drug may be modified in complex ways when drugs are combined. This establishes proof of concept that SEARCH-TB is a highly granular readout of drug interactions *in vivo,* providing information distinct from CFU burden and suggesting a future in which regimen selection is informed by *in vivo* molecular measures of *Mtb* physiology.

## INTRODUCTION

Tuberculosis (TB) is an ongoing public health crisis, causing ∼10 million illnesses and ∼1.4 million deaths each year.^1^ TB requires prolonged multi-drug treatment.^2^ To control TB, there is an urgent need for new regimens that cure both drug-susceptible and drug-resistant TB more quickly. Mouse treatment models play a crucial role in the preclinical evaluation of drugs and therapeutic regimens.^3–5^

In recent years, the number of promising individual new drugs and candidate compounds in the TB drug development pipeline has expanded.^6–8^ A key challenge is identifying combinations of these drugs that maximize efficacy. Unfortunately, conventional pharmacodynamic (PD) tools such as colony forming units (CFU) cannot reliably predict the efficacy of a drug combination based on the activity on individual drugs. Depending on their mechanism of action, drugs injure *Mycobacterium tuberculosis* (*Mtb*) differently, causing drug mechanism-specific physiologic perturbations.^9–11^ Since drug activity depends on the physiologic state of the bacterium,^12–14^ the effect of one drug may change the physiologic state of *Mtb* in a way that either favors the activity of a second antibiotic (synergy) or disfavors the activity of a second antibiotic (antagonism).

A limitation compounding understanding and prediction of combinatorial effects is that drug interactions are generally classified based on crude, culture-based measures of bacterial burden. Minimum inhibitory concentration (MIC) and minimum bactericidal concentration (MBC)^11^ estimate the threshold concentration at which bacterial burden remains constant or decreases 99%, respectively. Similarly, CFU, the PD marker conventionally used in animal studies, enumerates *Mtb* burden. More than 70 years ago, Eagle hypothesized that bacteria that are not immediately killed by drugs sustain damage to essential physiologic processes,^15^ initiating a “cascade of injury” that augments over time as homeostatic mechanisms are progressively disrupted. *Mtb* is also capable of adapting physiologic processes to withstand stresses, including drug exposure. PD markers of bacterial burden (*e.g.,* CFU) indicate only the presence or absence of the *Mtb,* providing no information about the complex processes of injury and adaption that is initiated by drug binding with its molecular target.

An alternative PD marker that captures the effect of drugs on bacterial physiologic state is the *Mtb* transcriptome. The effect of antibiotics on *Mtb* transcription has been studied extensively in short-term *in vitro* experiments,^16–22^ but short-term exposure in axenic culture may not replicate the physicochemical conditions and dynamic pharmacokinetics encountered during chronic *in vivo* exposure. To characterize drug effects *in vivo,* we recently developed SEARCH-TB, a targeted RNA-sequencing (RNA-seq) platform.^9,23^ Via a combination of selective eukaryotic cell lysis to deplete host RNA followed by targeted *Mtb* mRNA amplification, SEARCH-TB has enabled quantification of low-abundance *Mtb* transcripts in murine drug treatment models. Our evaluation of SEARCH-TB as a practical PD readout for preclinical drug and regimen evaluation began by evaluating the standard isoniazid, rifampin, pyrazinamide, ethambutol (HRZE) four-drug regimen^23^ and drugs in monotherapy.^9^ Here, we evaluate the capacity of SEARCH-TB to elucidate drug interactions in the BALB/c mouse model.

Historically, TB regimens have been selected based on a “mixing and matching” process in which the effect of iterative individual drug substitutions on burden or relapse outcomes is observed and compared. This empirical process led to the existing global standard 4-drug regimen for drug-susceptible TB (HRZE)^24^ and is used in contemporary human trials and murine studies. Unfortunately, this approach is time-consuming and can evaluate only a small fraction of possible drug/dose combinations.^25^ To move from empirical mix and match testing to rational regimen design, there is an unmet need for a PD marker that provides information about the effect of drug interactions on *Mtb* physiologic state *in vivo*. Our long-term goal is to develop SEARCH-TB as a PD tool for highly granular assessment of drug and regimen effects *in vivo*.

In the current work, we used SEARCH-TB to evaluate interactions among drugs in the bedaquiline, pretomanid, linezolid (BPaL) regimen, which recently revolutionized treatment of drug-resistant TB.^26,27^ Conducting a time-course study in the BALB/c mouse treatment model, which is a preclinical reference standard,^28^ we compared the effect of drugs individually, pairwise, and as a 3-drug combination on the physiologic state of *Mtb.* The novel SEARCH-TB *in vivo* physiologic readout revealed that drugs with different mechanism of action resulted in a distinct pattern of injury and adaptation that evolved with increasing treatment duration. In pairwise interactions, the effect of bedaquiline dominated interactions. In the full 3-drug regimen, the combination of pretomanid and linezolid made a transcriptionally synergistic contribution to bedaquiline.

## MATERIALS AND METHODS

### Murine Experiment and Sampling

We used RNA samples from a recent publication^29^ that fully describes murine procedures. Briefly, using the BALB/c high-dose aerosol infection model, mice were treated via oral gavage with standard doses of individual drugs, pairwise combinations, and the BPaL regimen (**Supplemental Table 1**). Mice were humanely euthanized before treatment initiation (pretreatment control, N = 5) or following 2, 4, 7, 11, 14, or 21 days of treatment (N = 5 per treatment/timepoint). CFU was quantified and RNA was preserved and extracted as previously described.^29^ All animal studies were performed at Colorado State University in a certified animal biosafety level III facility and conducted in accordance with guidelines of the Colorado State University Institutional Animal Care and Use Committee (reference number: 1212).

### Library Preparation and RNA sequencing

We prepared and sequenced RNA from mice euthanized before treatment initiation (pretreatment control) and 2, 7, and 14 days after treatment initiation. Samples were randomized and libraries were prepared in five different batches. We followed sequencing and quality control methods previously developed for the SEARCH-TB platform.^23^ Briefly, RNA was reverse transcribed, cDNA targets were amplified using the custom SEARCH-TB panel and sequencing was performed using an Illumina NovaSeq6000 device. The sequencing data was processed and quality control measures taken using pipelines previously described.^23^

### Statistical Analysis

To broadly characterize the differences in expression profiles between conditions and timepoints, we used principal components analysis. First, data was transformed using the variance stabilizing transformation (VST) from the DESeq2 package.^30^ To account for library preparation batch effect, we applied the removeBatchEffect function from the limma package to the normalized data.^31^ Principal components were then calculated using the 500 most variable genes.

We performed differential expression analysis by fitting negative binomial generalized linear models using the edgeR package.^32^ Models were fit using a term for drug condition/timepoint and a term for batch was also included to adjust for batch effect. Likelihood ratio tests were used to assess differences in expression between different conditions at each timepoint. Genes with Benjamini-Hochberg adjusted *p*-values^33^ of less than 0.05 for tests of interest were deemed significantly differentially expressed.

We performed hierarchical clustering of VST normalized data, averaged across samples in each condition/timepoint, to identify groups of genes with similar patterns of expression across time in the monotherapies. For each of the three monotherapies, we first removed invariant genes that were not differentially expressed between any timepoints and then performed hierarchical clustering using Euclidean distance and Ward’s method.^34^ Similar analysis was used to identify groups of genes that had differing expression patterns over time across the full BPaL regimen compared to bedaquiline monotherapy. For this analysis, we removed genes that showed no differential expression between bedaquiline and BPaL at any timepoint.

We performed enrichment analysis to identify functional gene categories that were enriched for genes significantly differentially expressed for comparisons of interest or for genes in clusters identified using hierarchical clustering. We used functional categories established by Cole et al.^35^ and curated from the literature (**Supplemental Table 2**), excluding categories with three or fewer genes. Enrichment was assessed using the hypergeometric test in the hypeR package^36^ with the test run separately for differentially expressed genes that were significantly upregulated and downregulated. Significance was assessed using a Benjamini-Hochberg adjusted *p*-value threshold of 0.05. We visualized the expression patterns for functional gene categories, as well as gene sets identified using hierarchical clustering, using sample-specific, scaled VST normalized expression values averaged across the genes in each cluster.

All analyses were performed using R (v4.4.1). Differential expression, functional enrichment, and visualizations can be explored interactively using an Online Analysis Tool [https://microbialmetrics.org/analysis-tools/].

## RESULTS

### Effect of individual drugs and combinations on CFU

Given as monotherapy, bedaquiline was bactericidal, decreasing median CFU by 99.2% (2.1 log10 difference) from pretreatment control to day 7 and 99.999% (5.17 log10 difference) by day 21 (**Fig 1a**). Pretomanid reduced CFU more gradually, by 27.78% (0.14 log10 difference) by day 7 and 99.49% (2.29 log10 difference) by day 21. Linezolid had a diminutive effect, reducing CFU 33.33% (0.18 log10 difference) by day 7 and 70% (0.52 log10 difference) by day 21. In pairwise combinations, linezolid did not discernably add to the bactericidal effect of either bedaquiline or pretomanid. As previously observed,^29,37^ the combination of pretomanid and bedaquiline had significantly less bactericidal activity than bedaquiline alone (Fig 1b, p-values <0.01 at days 7, 11, 14 and 21). Similarly, the BPaL regimen had significantly less effect on CFU than bedaquiline alone on days 7, 11 and 14 (p-values <0.003) although the 3-drug regimen and monotherapy were indistinguishable at day 21 (p-value=0.549) (**Fig 1b**).

**Figure 1.**
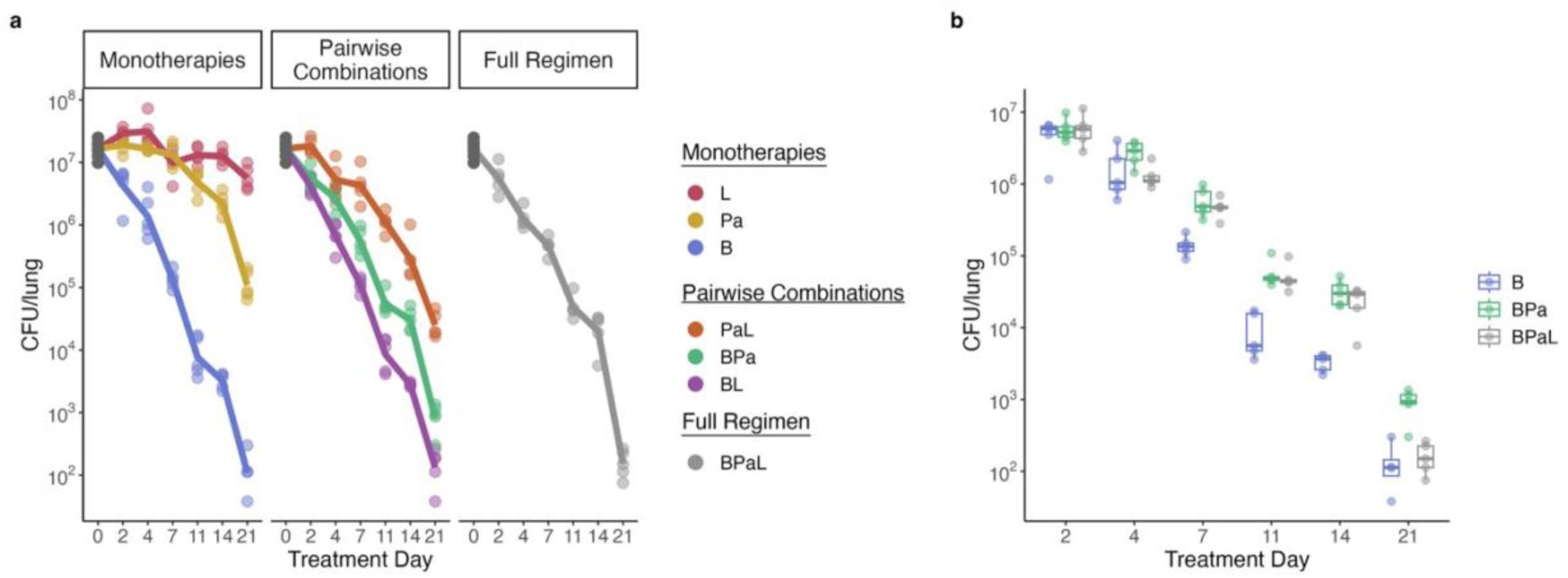
**Effect of individual drugs and combinations on CFU**. **a)** CFU burden over time in the lungs of BALB/c mice after treatment with bedaquiline (B), pretomanid (Pa), linezolid (L), and all combinations of the antibiotics. Points indicate CFU values from individual mice. For each treatment condition, lines connect the average value at each timepoint. **b)** Direct comparison of CFU for bedaquiline (B), BPa, and BPaL across time.

### Cascaded of injury and adaptation initiated by individual drug exposures

Monotherapy with bedaquiline, pretomanid or linezolid induced a rapid and profound transcriptional response, significantly altering the expression of 2,161 (60.6%), 1,304 (36.5%), and 947 (26.5%), *Mtb* genes respectively relative to the pretreatment control after two days of dosing (**Fig 2a**). With continued treatment, the *Mtb* transcriptional response to drug injury evolved further. For example, hierarchical clustering of differentially expressed genes revealed that bedaquiline exposure caused distinct sets of genes to change concordantly over time (**Fig 2b**). Cluster 1 consisted of 500 genes that had initially stable expression at day 2 followed by decreased expression at days 7 and 14 and was notably significantly enriched for ESX loci involved in active manipulation of host response (**Supplemental File 1**). Cluster 2 consisted of 887 genes with progressively decreased expression and was enriched for gene categories involved in translation, cell wall synthesis, and metabolism, indicating that bedaquiline-induced ATP starvation caused a global slow-down in cellular activity. Clusters 4 (443 genes) and 5 (708 genes) had progressively increased expression over time. In addition to generalized stress responses including insertion sequences and phage-related function, Cluster 5 was enriched for genes involved in regulatory function and specific responses which we have previously observed during adaptation to drug stress,^9^ including ESX4, Mce3 and fumarate reductase genes. Cluster 3 contained 199 genes that had decreased expression on day 2 and increased thereafter while cluster 6 contained 138 genes that increased on day 2 and decreased thereafter. Cluster 6 was enriched for transcription factors and other regulatory elements, consistent with a transient reprogramming with the onset of drug stress. Linezolid and pretomanid also caused progressive transcriptional change as illustrated in **Supplemental Figures S1** and **Supplemental Files 2-3**.

**Figure 2.**
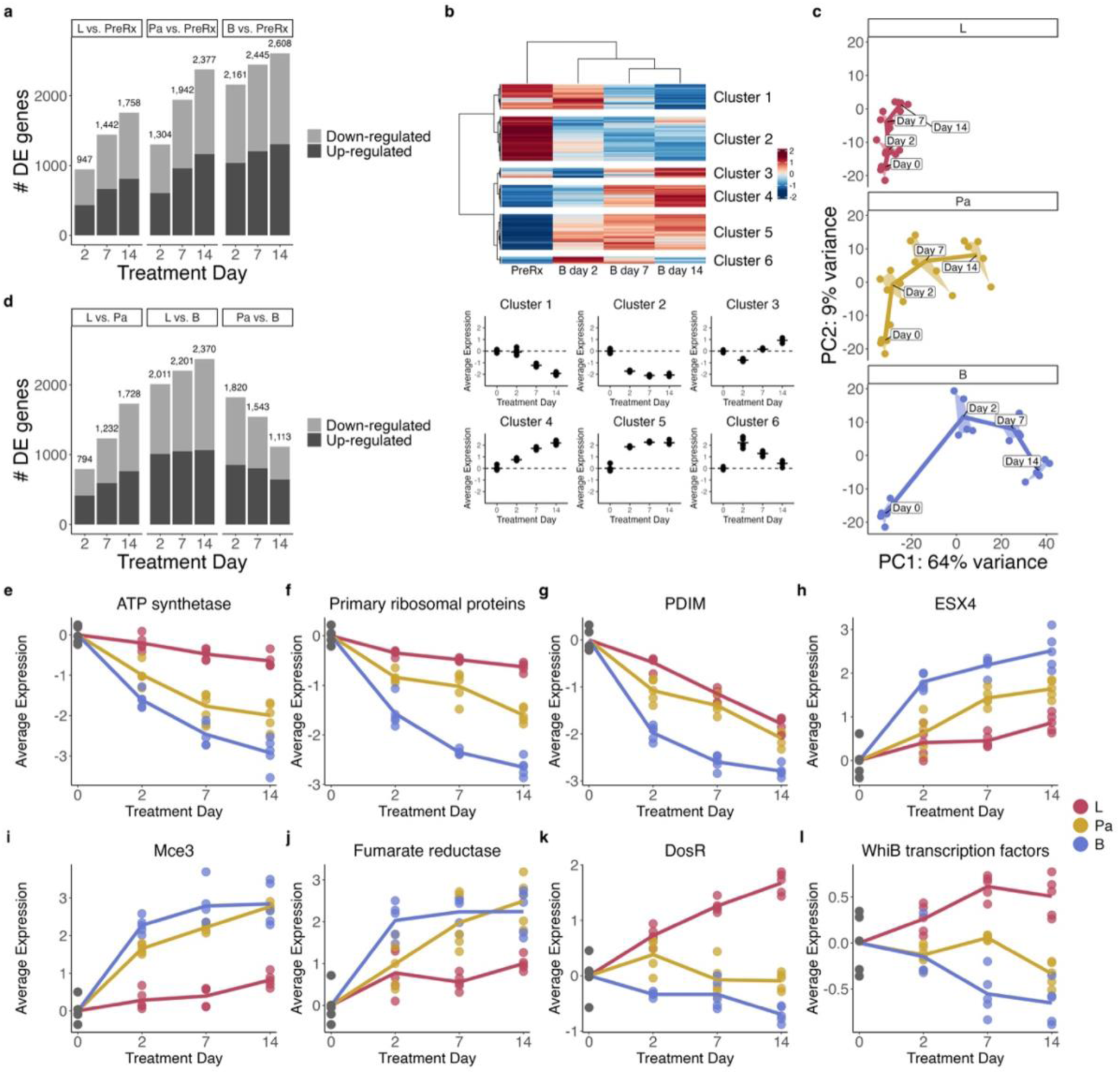
Effect of monotherapies on gene expression. **a)** Number of differentially expressed genes for bedaquiline (B), pretomanid (Pa), or linezolid (L) compared to pretreatment control at each timepoint. **b)** Hierarchical clusters of genes differentially expressed over time with bedaquiline treatment. Heatmap shows the VST normalized scaled gene expression averaged across samples. Clustering identified six broad patterns. Dot plots show the average expression for each cluster across timepoints. Each point represents an individual mouse. Horizontal lines indicate average values. Values are centered around the average value for the pretreated samples so that points above and below zero represent upregulation and downregulation relative to pretreatment, respectively. **c)** The first two principal components of VST-normalized gene expression data for the top 500 most variable genes for mice treated with each monotherapy across time. Each point represents an individual sample, and a convex hull highlights each timepoint. Timepoints are connected through the center of each convex hull. **d)** Number of differentially expressed genes between each pair of the individual antibiotics at each timepoint. **e) – l)** Average of VST-normalized, scaled gene expression in each monotherapy across time for genes in key Mtb biological processes: **e)** ATP synthetase, **f)** Primary ribosomal proteins, **g)** PDIM, **h)** ESX4, **i)** Mce3, **j)** fumarate reductase, **k)** DosR, **l)** WhiB transcription factors. Each point represents an individual mouse, and lines connect the mean for each timepoint. Values are centered around the average value for the pretreated samples so that points above and below zero represent upregulation and downregulation relative to untreated, respectively.

As we previously observed with first-line TB drugs,^9^ the patterns of injury and adaptation induced by bedaquiline, pretomanid, and linezolid were distinct both in terms of the magnitude and pattern of expression change. Principal Component Analysis (PCA) plots indicate that each drug led to a different *Mtb* transcriptional trajectory (**Fig 2c**). Direct comparison between drug-stressed phenotypes identified large numbers of differentially expressed genes (**Fig 2d**). Over time, the effects of bedaquiline and pretomanid grew progressively more distinct from linezolid.

The number of differentially expressed genes relative to linezolid increased from 2,011 to 2,370 for bedaquiline and from 794 to 1,728 for pretomanid between day 2 and day 14, respectively. Conversely, over time, the effects of bedaquiline and pretomanid grew more similar with the number of differentially expressed genes decreasing from 1,820 to 1,113 from day 2 to day 14.

Differences between the effects of bedaquiline, pretomanid, and linezolid on *Mtb* biological processes are too expansive to summarize comprehensively here so we describe only several examples. *Mtb* activity was inhibited most strongly by bedaquiline, followed by pretomanid, and then linezolid as highlighted by expression of genes coding for ATP synthetase, primary ribosomal proteins, and cell wall phthiocerol dimycocerosate (PDIM) (**Fig 2e-g**). Conversely, certain potentially adaptive responses, including genes coding for ESX4, Mce3, and fumarate reductase, increased most significantly with bedaquiline, followed by pretomanid and then linezolid (**Fig 2h-j**). Finally, there were gene categories with expression that changed divergently for linezolid compared to bedaquiline and pretomanid, including the DosR regulon, which responds to inhibited aerobic respiration (**Fig 2k**), and WhiB transcription factors, indicating a distinct regulatory response (**Fig 2l**). Biological processes changed by individual drugs can be explored via the Online Analysis Tool [https://microbialmetrics.org/analysis-tools/].

### Effect of pairwise drug combinations relative to the effect of monotherapy

#### Combination of bedaquiline and linezolid (BL)

The transcriptional perturbation caused by BL was similar to the effect of bedaquiline alone, as evidenced by the similar number of genes differentially expressed relative to pretreatment control (**Fig 3a**), the small number of genes differentially expressed in direct comparison between BL and bedaquiline alone (max = 85 genes differentially expressed at day 14) (**Fig 3b**), and the similar transcriptional trajectory in principal components over time between bedaquiline and BL (**Fig 3c**). By contrast, the effect of BL was highly distinct from the effect of linezolid alone, indicating the dominance of bedaquiline in the combination. This was shown by a substantially larger number of genes differentially expressed relative to pretreatment control for BL than for linezolid (**Fig 3a**), a large number of genes differentially expressed in direct comparison between BL and linezolid (**Fig 3b**) and distinct transcriptional trajectories in the principal components of linezolid compared with BL (**Fig 3c**).

**Figure 3.**
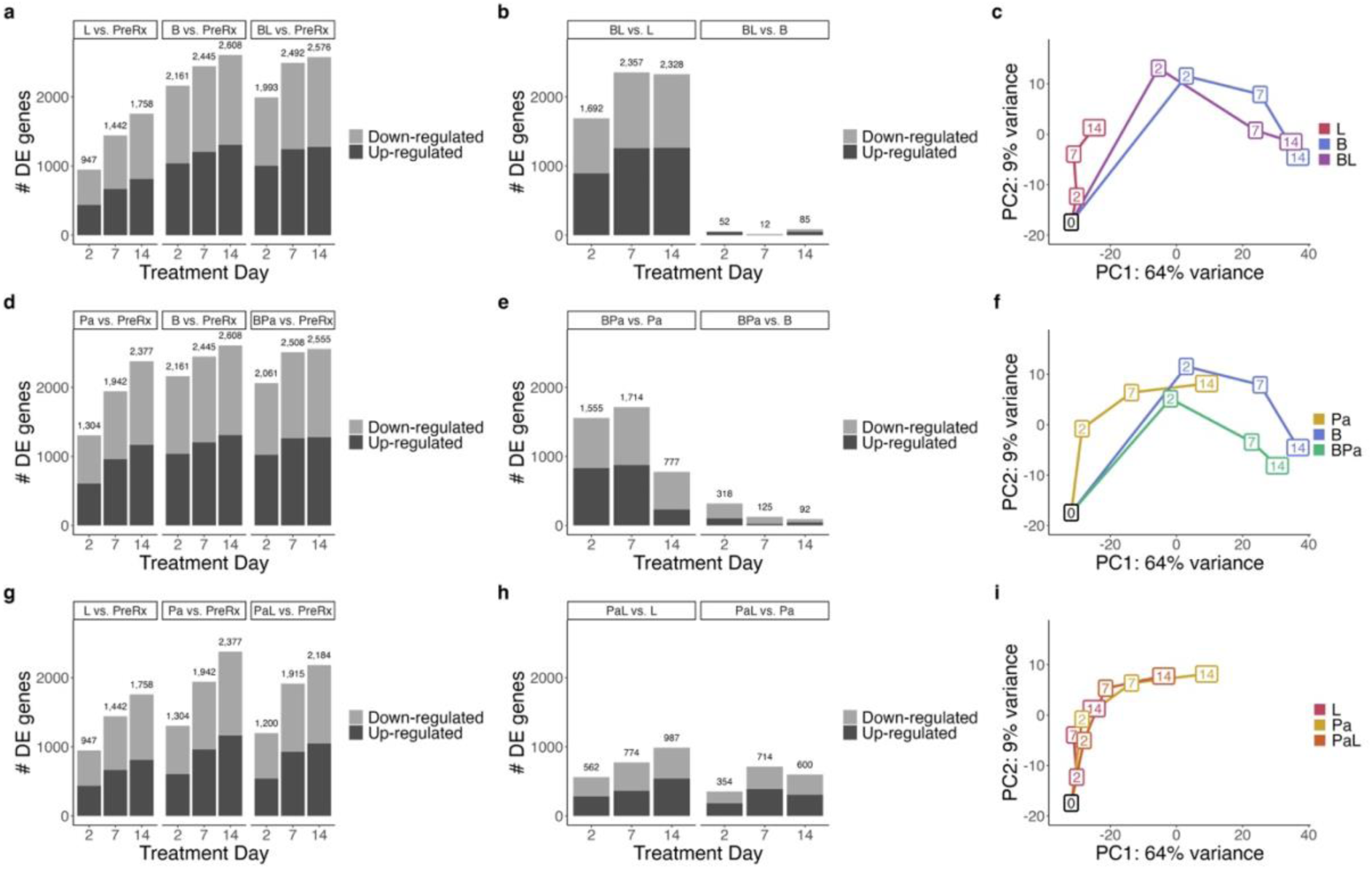
Effect of pairwise drug combinations relative to the effect of monotherapy. a)-c) Comparison of gene expression results for bedaquiline (B), linezolid (L), and the combination (BL). **d)-f)** Comparison of gene expression results for bedaquiline (B), pretomanid (Pa), and the combination (BPa). **g)-i)** Comparison of gene expression results for linezolid (L), pretomanid (Pa), and the combination (PaL). **a), d)** and **g)** show the number of differentially expressed genes for the two individual antibiotics and their pairwise combination relative to the pretreatment condition. **b), e)** and **h)** show the number of differentially expressed genes for the pairwise combination relative to the individual monotherapies. **c), f)** and **i)** show first two principal components of VST-normalized gene expression data for the top 500 most variable genes for the two individual antibiotics and their pairwise combination. Each box represents the average of the principal components across timepoint/treatment with the number in each box representing the treatment day. Colors of the boxes/lines represent different treatments.

#### Combination of bedaquiline and pretomanid (BPa)

The transcriptional change caused by BPa differed modestly from the effect of bedaquiline alone. The number of genes differentially expressed relative to pretreatment control was similar for BPa and bedaquiline alone (**Fig 3d**). Following day 2, at which point 316 genes were significant, relatively few genes were differentially expressed in direct comparison between BPa and bedaquiline (**Fig 3e**). BPa and bedaquiline had similar transcriptional trajectories in PCA analysis (**Fig 3f**). By contrast, the effect of BPa was highly distinct from the effect of pretomanid alone, again highlighting the dominance of bedaquiline in the combination. This was shown by the substantially larger number of genes differentially expressed relative to pretreatment control for BPa than for pretomanid alone (**Fig 3d**), a large number of genes differentially expressed in direct comparison between BPa and pretomanid (**Fig 3e**) and the distinct transcriptional trajectories in PCA analysis (**Fig 3f**).

#### Combination of pretomanid and linezolid (PaL)

The transcriptional change induced by PaL was distinct from the effect of either pretomanid or linezolid alone (**Fig 3g-i)**. For example, on day 14, pretomanid and linezolid had 600 and 987 differentially expressed genes relative to PaL, respectively (**Fig 3h**). The transcriptional trajectory of PaL was intermediate between either pretomanid or linezolid, indicating a blended effect.

### Effect of BPaL relative to bedaquiline alone

Because preceding analyses indicated that bedaquiline dominates pairwise combinations, we evaluated the effect of adding both pretomanid and linezolid to bedaquiline. We found that PaL modified the effect of bedaquiline synergistically in the sense that the number of genes significantly differentially expressed between BPaL and bedaquiline was greater than expected based on the addition of either drug alone. For example, at day 14, BPa and BL differed from bedaquiline by 92 and 85 genes respectively but BPaL differed from bedaquiline by 500 genes (**Fig 4a**). Transcriptional synergy was also observed at days 2 and 7. PCA indicates that BPaL led to a distinct longitudinal trajectory relative to bedaquiline alone (**Fig 4b**).

**Figure 4.**
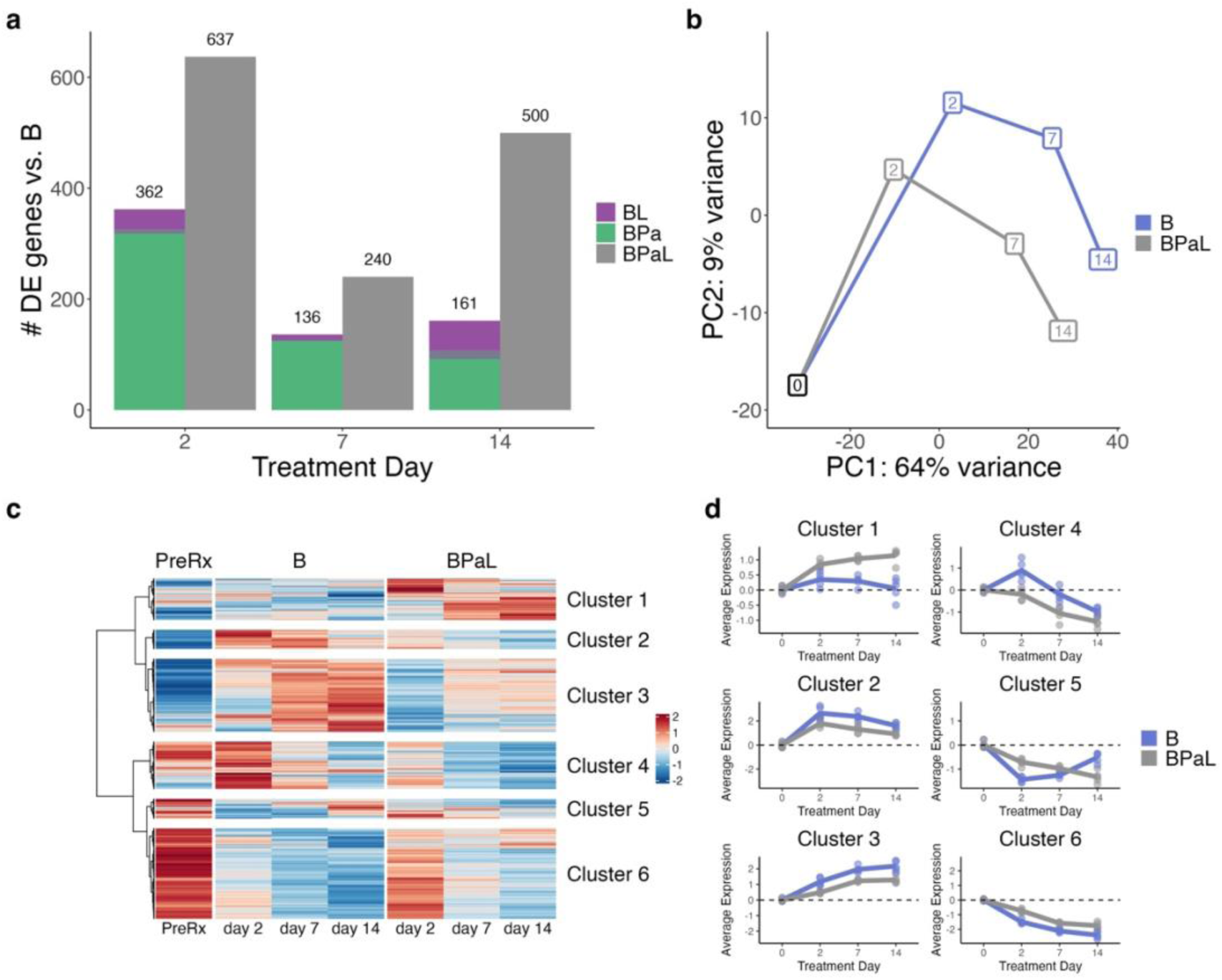
Effect of BPaL relative to bedaquiline alone. **a)** Number of differentially expressed genes for the pairwise combinations containing bedaquiline (BL and BPa) relative to bedaquiline (B) alone, compared to the number of differentially expressed genes of the full BPaL regimen relative to bedaquiline alone. **b)** The first two principal components of VST-normalized gene expression data for the top 500 most variable genes for B and the BPaL regimen. Each box represents the average of the principal components across timepoint/treatment with the number in each box representing the treatment day. Colors of the boxes/lines represent the treatment. **c)** Heatmap showing the VST normalized scaled gene expression averaged across samples for genes differentially expressed between B and BPaL at any timepoint. Hierarchical clustering identified six broad patterns. **d)** Average expression for each cluster from plot (c) across timepoints for B and BPaL samples. Each point represents an individual mouse. Lines connect the average value for each treatment across timepoints. Points and lines are colored by treatment. Values are centered around the average value for the pretreated samples so that points above and below zero represent upregulation and downregulation relative to pretreatment, respectively. Plots for each cluster have unique scales to better show the differences between B and BPaL

To elucidate the differing effects of BPaL relative to bedaquiline monotherapy, we evaluated longitudinal change of genes differentially expressed at any timepoint between BPaL and bedaquiline monotherapy and identified six gene clusters with distinct patterns of change across the two therapies (**Fig 4c-d**). Cluster 1 contained genes induced to a greater degree by BPaL than by bedaquiline monotherapy but was not significantly enriched for any gene sets (**Supplemental File 4**). Conversely, Clusters 2 and 3 comprised genes that were induced to a lesser degree by BPaL than by bedaquiline monotherapy. Cluster 2 was statistically enriched for genes involved in regulatory functions, suggesting transient reprogramming in response to initial drug stress.

Cluster 3 was statistically enriched for phage-related functions, a non-specific stress response. Cluster 4 contained genes that were initially induced in bedaquiline alone, before falling across later timepoints, but gradually decreased in BPaL. Cluster 4 was statistically enriched for regulatory and stress responses. Conversely, Cluster 5 was transiently repressed on day 2 but then rose over time for bedaquiline monotherapy while continuing to decrease for BPaL. Cluster 5 was enriched for three categories related to cholesterol catabolism, suggesting BPaL caused sustained repression while bedaquiline allowed recovery of this important metabolic function.

Cluster 6 was repressed more rapidly and to a greater degree over time by bedaquiline monotherapy than by BPaL. This cluster was enriched for categories related to metabolism (TCA cycle, KstR1), protein synthesis (ribosomal proteins, protein translation and modification) and cell wall synthesis (PDIM, fatty acid synthases II), broadly indicating greater inhibition of activity by bedaquiline alone.

## DISCUSSION

SEARCH-TB is a novel candidate PD marker that evaluates the effect of drugs on *Mtb* physiologic processes rather than *Mtb* burden. Here we evaluated the capacity of SEARCH-TB to elucidate drug interactions in the BPaL regimen in a mouse TB treatment model. When given in monotherapy, each drug elicited a distinct transcriptional response that evolved over time, consistent with a cascade of progressive physiologic injury and adaptation. When given in pairwise combinations, bedaquiline dominated the effect of either pretomanid or linezolid.

When given as a 3-drug combination, pretomanid and linezolid appeared to modify the effect of bedaquiline synergistically, producing a transcriptional change greater than would be expected for either drug alone. The SEACH-TB molecular analysis of physiologic processes provided vastly more granular information than CFU alone. This work positions SEARCH-TB as a highly granular pharmacodynamic tool for understanding the effect of drugs and regimens *in vivo*.

These results confirmed and expanded upon our previous study of first-line drugs which identified drug-specific transcriptional changes after one month of treatment in the BALB/c mouse^9^ by performing a time-course study and evaluating two additional drugs (pretomanid and linezolid). For all three drugs, the *Mtb* transcriptional response grew progressively greater in number and fold-change compared to the pretreatment control, suggesting progressive loss of homeostasis that presumably culminates in cell death. Particularly notable was the scale and rapidity of *Mtb* transcriptional change. For example, bedaquiline at a human-equivalent dose significantly altered expression of 61% of genes assayed after only two days. Down-regulation of genes related to macromolecule synthesis, metabolism, and growth suggested that inhibition of ATP synthesis rapidly incapacitated *Mtb*. There were both broad consistencies and important differences between the days-long response to bedaquiline we observed in this study in mice and responses observed in previous short *in vitro* experiments.^38^ For example, bedaquiline-mediated inhibition of ATP synthesis suppressed *Mtb* metabolism and macromolecule synthesis *in vitro* and *in vivo*, but certain adaptations such as expression of the DosR regulon changed divergently, increasing significantly *in vitro* and decreasing significantly *in vivo.* Interestingly, pretomanid appeared to “follow” bedaquiline over time such that the day 14 pretomanid transcriptome was similar to the day 2 bedaquiline transcriptome. By contrast, the linezolid transcriptome progressively diverged from both pretomanid and bedaquiline over time. These monotherapy time-course exposure results indicate that each drug’s initial engagement with its molecular target initiates distinct secondary cascades of injury and adaptation. Finally, the monotherapy results highlight that SEARCH-TB reveals drug effects that are indiscernible based on CFU. In particular, CFU showed that linezolid had a static effect, suggesting minimal activity. By contrast, SEARCH-TB showed that linezolid altered expression of thousands of genes, indicating a shift in *Mtb* physiology.

SEARCH-TB for pairwise combinations showed that BL or BPa closely resembled bedaquiline alone. The dominance of the bedaquiline effect is consistent with the central role of bedaquiline in contemporary treatment regimens. By contrast, the combination of pretomanid and linezolid produced a transcriptional response that was intermediate between either drug, demonstrating that each drug modifies the pattern of injury and adaptation induced by the other.

Whereas addition of a single drug to bedaquiline had a small effect, addition of both pretomanid and linezolid to bedaquiline altered the *Mtb* transcriptome more than anticipated based on the results of addition of either pretomanid or linezolid to bedaquiline individually. Addition of PaL appeared to mitigate the bedaquiline-induced suppression of metabolism, protein synthesis, and cell wall synthesis, indicating less suppression of *Mtb* activity with BPaL. Since a more active phenotype might be more capable of recovery on agar, this may explain the seemingly paradoxical but well-established^29,37^ finding that BPaL decreased CFU counts less than bedaquiline alone. In contrast, other processes exhibited a greater change at the end of treatment with BPaL than with bedaquiline alone. For example, gene sets associated with stress response were more repressed by BPaL than by bedaquiline alone. Additionally, a gene set that decreased more for BPaL than for bedaquiline alone (Cluster 5) was enriched for several cholesterol pathways, suggesting that the addition of PaL further inhibited cholesterol catabolism. Cholesterol catabolism is essential for *Mtb* intracellular survival^39,40^ and is a promising novel drug target.^2^ Others have previously shown that the cholesterol inhibitor GSK- 286 enhanced efficacy to a greater degree when added to BPaL than when added to BPa.^41^ This highlights a common observation in murine studies: the effect of adding a single drug to a regimen often depends on the specific combination of drugs it is added to.^42–44^ We hypothesize that the activity of one drug may be conditional on the pattern of injury and adaptation induced by other drugs in the regimen. By elucidating *Mtb* physiology *in vivo,* SEARCH-TB provides a novel basis for understanding these complex drug interactions.

This work adds to the body of evidence^9,23^ that SEARCH-TB is a highly granular and informative PD marker. CFU enumerates the burden of *Mtb* capable of growth on agar but provides no information on bacterial physiologic processes. By contrast, SEARCH-TB provides no information about *Mtb* burden but gives rich information about the cascade of injury and adaptation triggered by drug stress. Here we found that SEARCH-TB gave a markedly different perspective on drug interactions than CFU. For example, BPa and BPaL had indistinguishable effects on CFU before day 21. By contrast, SEARCH-TB showed that the addition of linezolid to BPa meaningfully altered *Mtb* physiology.

To date, our SEARCH-TB studies have been descriptive, documenting the effect of HRZE,^23^ individual drugs^9^ and now drug interactions in the BALB/c mouse. SEARCH-TB could provide practical value for drug evaluation if patterns of injury and adaptation measured by the transcriptome can predict drug activity and/or interactions *in vivo*. The INDIGO-MTB model provided proof of concept that transcriptional responses to drug exposure *in vitro* can predict drug combinations that optimally inhibit growth *in vivo.*^45^ A next step will be assessing the ability of SEARCH-TB to predict the treatment shortening (*i.e.,* sterilizing activity) of regimens by determining whether specific patterns of injury and adaptation are associated with faster time to non-relapsing cure in mice (*i.e.,* shorter T95 in the BALB/c relapsing mouse model).

Identifying an early signature of treatment shortening in the BALB/c model could reduce reliance on the long and resource intensive relapsing mouse model that is a bottleneck in regimen evaluation and initiate a new era of drug and regimen evaluation based on highly granular molecular measures of injury and adaptation rather than crude measures of culturable *Mtb* burden.

This study has several limitations. First, we selected the BALB/c high-dose aerosol infection model because it is the cornerstone of contemporary regimen development. Drug effects in other TB treatment models such as the C3HeB/FeJ mouse may differ due to variation in pharmacokinetics and tissue microenvironments. A future direction will be applying SEARCH-TB to evaluate bacterial phenotypes and drug responses in diverse tissue types. Second, as performed here, SEARCH-TB quantified the population-average bacterial transcriptome rather than capturing heterogeneity within the bacterial population. Future studies will evaluate the capacity of the highly sensitive SEARCH-TB method to evaluate gene expression within individual host cells. Finally, while the transcriptome documents the cascade of physiologic change initiated by drug stress, it cannot disentangle what components of this change are functional (*i.e.,* adaptations that enable survival) versus dysfunctional (*i.e.,* “damage” that results from profound physiologic stress).

Using SEARCH-TB allowed us to assess drug interactions within the BPaL regimen *in vivo* based on how they affect physiologic processes. This revealed that individual drugs initiate distinct cascades of injury and adaptation that are modified in complex ways when drugs are combined, resulting in either dominance by a single drug (*i.e.,* bedaquiline) or a blended response. The response to the full 3-drug BPaL regimen indicates that higher-order interactions may be more than the sum of parts. Broadly, this work demonstrates the capacity of SEARCH-TB to elucidate drug interactions with vastly greater granularity than the existing conventional measure of culturable *Mtb* burden (CFU), pointing to a future in which optimal combinations could be identified based on *in vivo* molecular measures of *Mtb* physiologic processes.

