## Supplemental Information for "Deconvoluting drug interactions based on *M. tuberculosis* physiologic processes: Transcriptional disaggregation of the BPaL regimen *in vivo*"

### **Supplemental Methods**

#### **Table S1.** Antibiotic dosing

Antibiotic dosage concentration for mice in each antibiotic treatment group. Mice began treatment 11 days post infection via oral gavage seven days a week.

| **Treatment** | **Dose (mg/kg)** |
| --- | --- |
| Bedaquiline | 25 |
| Linezolid | 100 |
| Pretomanid | 50 |

#### **Table S2.** Curated gene categories used for enrichment analysis.

Manuscript figures and statistical functional enrichment analysis used sets of biologically linked genes curated from the literature. The number of genes in category indicates the number of genes identified in the literature source. Since SEARCH-TB quantifies expression of 89% of *Mtb* transcripts, not all genes in each category were quantified. Number of genes in SEARCH-TB indicates the number assayed.

| **Category** | **# of Genes in SEARCH-TB** | **# of Genes in Category** | **Source** |
| --- | --- | --- | --- |
| ABC transporters - Type I peptide and amino acids | 6 | 8 | (Soni, Dubey, & Bhatnagar, 2020)^1^ |
| ABC transporters - Type I Sugar Import | 12 | 12 | (Soni, Dubey, & Bhatnagar, 2020)^1^ |
| ABC transporters - Type I anion | 7 | 7 | (Soni, Dubey, & Bhatnagar, 2020)^1^ |
| ABC transporters - Type I phosphate | 8 | 8 | (Soni, Dubey, & Bhatnagar, 2020)^1^ |
| ABC transporters - Type II metal | 5 | 5 | (Soni, Dubey, & Bhatnagar, 2020)^1^ |
| Alternative ribosomal proteins | 4 | 5 | (Prisic et al., 2015)^2^ |
| Antigen 85 | 3 | 3 | (Karbalaei Zadeh Babaki, Soleimanpour, & Rezaee, 2017)^3^ |
| Antitoxins | 72 | 76 | (Shao et al., 2011)^4^ |
| Arabinogalactan (AG) | 18 | 19 | (Abrahams & Besra, 2018)^5^ |
| Beta Oxidation | 18 | 18 | (Schnappinger et al., 2003)^6^ |
| Cell wall synthesis | 40 | 40 | (Kirksey et al., 2011)^7^ |
| Cholesterol A and B ring degradation | 10 | 10 | (Pawełczyk et al., 2021)^8^ |
| Cholesterol C and D ring degradation | 5 | 5 | (Pawełczyk et al., 2021)^8^ |
| Cholesterol side chain degradation | 33 | 33 | (Pawełczyk et al., 2021)^8^ |
| Cutinase-Like Proteins (CULP) | 7 | 7 | (Tallman, Levine, & Beatty, 2016)^9^ |
| Cytochrome oxidase bccaa3 | 6 | 7 | (Lee, Sviriaeva, & Pethe, 2020)^10^ |
| Cytochrome oxidase bd | 3 | 4 | (Lee, Sviriaeva, & Pethe, 2020)^10^ |
| DNA replication and repair | 25 | 27 | (Ditse, Lamers, & Warner, 2017)^11^ |
| DosR | 48 | 48 | (Voskuil et al., 2003)^12^ |
| Drug targets | 40 | 40 | (“Working Group for New TB Drugs.,” 2021)^13^  (Shetye, Franzblau, & Cho, 2020)^14^ |
| Efflux Pumps and Transports | 25 | 26 | (Remm, Earp, Dick, Dartois, & Seeger, 2022)^15^ |
| Enduring Hypoxic Response | 149 | 161 | (Rustad, Harrell, Liao, & Sherman, 2008)^16^ |
| Esterases (Lip family) | 20 | 22 | (Tallman, Levine, & Beatty, 2016)^9^ |
| Esterases (non-Lip family) | 13 | 13 | (Tallman, Levine, & Beatty, 2016)^9^ |
| ESX1 | 18 | 19 | (Gröschel, Sayes, Simeone, Majlessi, & Brosch, 2016)^17^ |
| ESX2 | 12 | 12 | (Gröschel, Sayes, Simeone, Majlessi, & Brosch, 2016)^17^ |
| ESX3 | 9 | 11 | (Gröschel, Sayes, Simeone, Majlessi, & Brosch, 2016)^17^ |
| ESX4 | 7 | 7 | (Gröschel, Sayes, Simeone, Majlessi, & Brosch, 2016)^17^ |
| ESX5 | 11 | 15 | (Gröschel, Sayes, Simeone, Majlessi, & Brosch, 2016)^17^ |
| Fatty Acid Synthases I | 1 | 1 | (Cole et al., 1998)^18^ |
| Fatty Acid Synthases II | 8 | 9 | (Duan, Xiang, & Xie, 2014)^19^ |
| Fumarate reductase | 4 | 4 | (Cole et al., 1998)^18^ |
| Kas operon | 3 | 5 | (Slayden & Barry, 2002)^20^ |
| kstR1 regulon | 70 | 71 | (Wipperman, Sampson, & Thomas, 2014)^21^ |
| kstR2 regulon | 14 | 15 | (Wipperman, Sampson, & Thomas, 2014)^21^ |
| LpqY-SugA-SugB-Sug trehalose transporter | 5 | 5 | (Soni, Dubey, & Bhatnagar, 2020)^1^ |
| Mce1 | 7 | 7 | (Cole et al., 1998)^18^ |
| Mce2 | 6 | 7 | (Cole et al., 1998)^18^ |
| Mce3 | 7 | 7 | (Cole et al., 1998)^18^ |
| Mce4 | 7 | 7 | (Cole et al., 1998)^18^ |
| LAM | 14 | 15 | (Batt, Burke, Moorey, & Besra, 2020)^22^ |
| mmpL | 14 | 14 | (Domenech, Reed, & Barry, 2005)^23^ |
| mmpS | 5 | 5 | (Melly & Purdy, 2019)^24^ |
| Mycobactin Biogenesis | 10 | 10 | (Quadri, Sello, Keating, Weinreb, & Walsh, 1998)^25^ |
| Mycolic acid condensation and transfer | 7 | 7 | (Marrakchi, Lanéelle, & Daffé, 2014)^26^ |
| Mycolic acid modification | 12 | 12 | (Marrakchi, Lanéelle, & Daffé, 2014)^26^ |
| NADH dehydrogenase type I | 12 | 14 | (Cook, Hards, Vilchèze, Hartman, & Berney, 2014)^27^ |
| NADH dehydrogenase type II | 2 | 2 | (Cook, Hards, Vilchèze, Hartman, & Berney, 2014)^27^ |
| Nitrate import and reductase | 6 | 8 | (Cole et al., 1998)^18^ |
| Oxidative Stress | 48 | 49 | (Voskuil, Bartek, Visconti, & Schoolnik, 2011)^28^ |
| PDIM | 20 | 20 | (Rens, Chao, Sexton, Tocheva, & Av-Gay, 2021)^29^ |
| Peptidoglycan (PG) | 32 | 34 | (Maitra et al., 2019)^30^ |
| Phospholipase C | 4 | 4 | (Raynaud et al., 2002)^31^ |
| Primary ribosomal proteins | 50 | 53 | (Prisic et al., 2015)^2^, (Cole et al., 1998)^18^* |
| Sigma Factors | 12 | 13 | (Lew, Kapopoulou, Jones, & Cole, 2011)^32^ |
| Stringent Response - Induced | 58 | 70 | (Dahl et al., 2003)^33^ |
| Stringent Response - Repressed | 66 | 78 | (Dahl et al., 2003)^33^ |
| Succinate dehydrogenase I and II | 7 | 7 | (Hartman et al., 2014)^34^ |
| Toxin-Antitoxin | 146 | 152 | (Shao et al., 2011)^4^ |
| Toxins | 74 | 76 | (Shao et al., 2011)^4^ |
| Transcription Factors | 187 | 198 | (Lew, Kapopoulou, Jones, & Cole, 2011)^32^ |
| Trehalose | 10 | 10 | (Wilson et al., 1999)^35^ |
| Triacylglycerol Synthases | 14 | 16 | (Thanna & Sucheck, 2016)^36^ |
| UgpABCE glycerophosphocholine transporter | 4 | 4 | (Soni, Dubey, & Bhatnagar, 2020)^1^ |
| Universal stress proteins | 9 | 10 | (Lew, Kapopoulou, Jones, & Cole, 2011)^32^ |
| UspABC amino sugar transporter | 3 | 3 | (Soni, Dubey, & Bhatnagar, 2020)^1^ |
| WhiB-Like transcription factors | 7 | 7 | (Wan et al., 2021)^37^ |
| Zur regulon | 17 | 20 | (Dow et al., 2021)^38^ |

* Primary ribosomal proteins category was created by removing the four alternative proteins identified in Prisic et al., 2015 from the “Ribosomal protein and synthesis” gene list identified in Cole et al., 1998.

**Supplemental Results**

#### **Figure S1:** Change in gene expression over time for Pa and L.

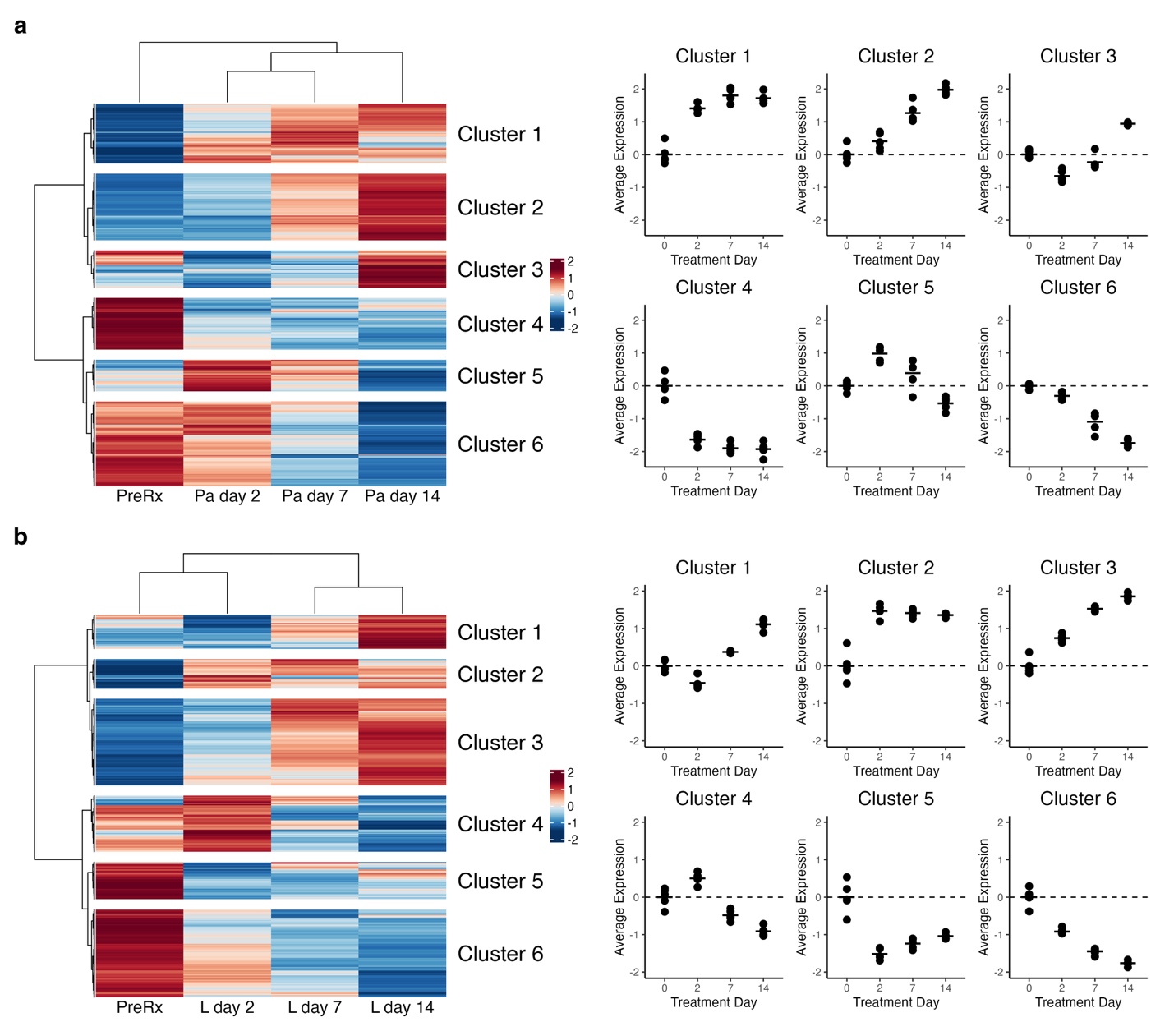

**Figure S1.** Change in gene expression over time for Pa (a) and L (b)**.** Heatmaps show the VST normalized scaled gene expression averaged across samples. Genes that were not differentially expressed between at least two timepoints were excluded. For each monotherapy, hierarchical clustering identified six broad patterns. Dot plots show the average of VST-normalized, scaled expression across timepoints for the clusters. Each point represents an individual mouse. Horizontal lines indicate average values. Values are centered around the average value for the pretreated samples so that points above and below zero represent upregulation and downregulation relative to pretreatment, respectively.
